# The effect of grey wolf (*Canis lupus*) and human disturbance on the activity of big game species in the Bükk Hills, Hungary

**DOI:** 10.1101/2022.09.21.508874

**Authors:** Zsófia Szabó, Péter Gombkötő, Sándor Csaba Aranyi, László Patkó, Dóra Gigler, Zoltán Barta

## Abstract

The recent return of wolves to the Hungarian forests escalates conflicts among stakeholders. Hunting management agencies communicate that the presence of wolves may change the behaviour of big game species leading to difficulties for hunting organization and logistics. Here, we take a data driven approach to explore the activity of wolves and big game species. For this purpose we analysed camera trap data, collected in the Bükk National Park, Hungary. To estimate avoidance among wolves, humans and games we calculated a non-parametric activity overlap coefficient (Δ_4_) and used a machine learning (XGBoost) model. Our results show that game species have higher overlap coefficient with wolf (Δ_4_ = 0.83-0.89) than with human activity (Δ_4_ = 0.26-0.52), because predators and games are active in the same periods of the day, mainly night and dawn, and human activity mainly takes place during daytime. We could detect the refugee effects in the case of all game species. Our XGBoost analyses only found a moderate negative effect of wolf on red deer occurrence, while human activity had higher importance value and lowered the occurrence of all three game species investigated. Our results may thus indicate that human disturbance might be more important in shaping game activity than the presence of the grey wolf in Hungary.

## Introduction

Earlier studies suggest that grey wolf can have an important role in the ecosystem as a top predator. For example, the processes initiated by the reintroduction of wolves in the Yellowstone National Park in 1995-1996 supports the Trophic Cascade Theory. After the wolf reintroduction, the elk (*Cervus elaphus*) population decreased and their spatial distribution was altered. Due to these changes the vegetation could have renewed and provided habitat and food for many other species. Consequently, the elks population without wolves seemed to have an increased browsing effect, which led to species decline, while it appears that the wolves indirectly contributed to the growth of biodiversity by their direct and indirect effects on elks [1–4]. The significance of trophic cascades caused by large carnivores, however, still remains questionable in many other cases [5,6].

But not only carnivores can influence game populations. Nowadays, constant human presence is also widespread in many of the forests. The pioneering studies of Hediger (1934) (cited in [7]) and Walther (1969) about fleeing distances have initiated research on the effect of human disturbance on African game species. According to their results, games produce the same anti-predator response to approaching cars as to predators. Since then, a lot of research has supported [8–10] that this anti-predator behaviour is not only triggered by lethal human disturbance (e.g. hunting) but also by non-lethal ones (e.g. hiking, [11]). Nevertheless, it seems that games do not indiscriminately respond to different disturbances, their responses depend on the type of disturbance [9–11]. For example, the games may respond less intensively to a hiker or horse rider than to logging or hunting [9,11]. However, according to a meta-analysis by Doherty et al. (2021) [12], hunters and hikers may have a larger effect on the movements of animals than urbanisation or logging. A form of these anti-predator behaviour can be the avoidance of disturbing agents though limiting the spatial and temporal overlap with them [13–16].

Behaviourally mediated direct effects can be observed between predator and game species when the game’s avoidance of predators has an influence on the distribution of games [17,18]. In the same way humans can influence the distribution of predators [1,8,19], but this does not mean that humans do not influence the distribution of game individuals. Hebblewhite et al. (2005) [1] suggests that elks would avoid areas where human disturbance is low, but the abundance of wolves is high; an hypothesis is supported by several other studies, e.g. [10,14,15]. The refugee effect is, when an area with high human disturbance may serve as refugees for preys, because predators avoid humans more than their preys do [15,20,21]. So whether ungulates avoid wolf-dens areas and prefer human dominated landscapes depends on the type of human land use [9]. Moreover, based on a meta-analysis by Tucker et al. 2022 [22], animal movement activity decreases where human impact is high.

Games can also adapt to predators and human disturbance by limiting their temporal overlap in their activities. According to previous observations, games often change their daily routines if a predator appears in the same habitat [23–27]. Predators, however, are able to adjust their activities to match them to the games’ [27,28]. Also human disturbance can change the predator’s activity patterns indirectly through the changes caused in activity patterns of games [28].

The grey wolf (*Canis lupus*) is a native species in Hungary; its extinction in the late 19th century was solely caused by human persecution and extermination [29–31]. Until that time its presence in the Carpathian Basin was continuous for at least the last 15 000-16 000 years in spite of extensive climate changes [30]. Their sporadic occurrence has been documented again since the 1970s in the northeast part of Hungary, near Slovakia [32]. The presence of wolves in the territory of Bükk National Park has been reported in 2010 by Bükk National Park Directorate [33]. Now, Hungary has stable populations in the northeast part of the country, in the Bükk Region (25-35 ind., [34]), Aggtelek Karst (12 ind.) and Zemplén Mountains (25 ind.) of the North Hungarian Mountains [35].

According to a study carried out in the Aggtelek National Park [36] the main food of wolves were big game, such as red deer (*Cervus elaphus*) and wild boar (*Sus scrofa*) followed by roe deer (*Capreolus capreolus*) and muflon (*Ovis gmelini*). From the point of view of nature conservation it is welcomed that wolves take control of the population size of game species, but the hunting industry became worried that the presence of wolves will reduce their efficiency in big game hunting [37]. For instance, they communicate that hunting has become more difficult due to the big game’s behavioural changes caused by wolves: games started to avoid certain areas and became more shy [38–41].

Currently, both advantageous and disadvantageous effects of wolves for certain stakeholder groups are, however, only supported by anecdotal evidence in Hungary. Furthermore, the arguments in general do not take into account that human presence might be as, or even more, crucial for game as of the predators. Here, we aim to contribute to this debate in an evidence based way, by quantitatively analysing camera trap data, recorded in the Bükk National Park, to investigate the avoidance by machine learning and the temporal overlap between activities of games and predators and human disturbance. We assume that negative associations between game and either predator or human disturbance is an indication that game avoids predators or humans.

## Materials and Methods

### Study area

Our research was carried out in the central area of the Bükk Mountains, in the Bükk National Park (BNP), northern Hungary (N 48°06’, E 20°30’). The largest forested area of Hungary (*c*. 140 km^2^) can also be found here. This is one of the coldest parts of Hungary with mean annual temperature varying between 7-8 °C. The mean yearly rainfall, 600-700 mm, is high compared to the rest of the country [42]. The number of days with snow cover is also outstanding within Hungary [42,43]. Forest cover is 95% formed by species like mountain and submountain beech (*Fagus sylvatica*), sessile oak (*Quercus patreae*) and Austrian oak (*Quercus cerris*) [42–44]. The main economic activity is forestry, managed by two large state-owned companies (Egererdő Forestry Company and Északerdő Forestry Company). Many roads cross the study area, and it is a popular place for hunting, hiking and other recreational activities. Vehicular traffic consists of off-road vehicles, cars, trucks, tractors and motorcycles. Hikers mainly move on foot or ride on horse or bicycle. Seasonally there are many mushroom collectors as well (pers. obs. P. Gombkötő).

The big game species in the area (with their estimated numbers after Csányi 2022 [45] are red deer (*Cervus elaphus*, 10 085), boar (*Sus scrofa*, 2 465), muflon (*Ovis gmelini*, 2 332) and roe deer (*Capreolus capreolus*, 10 741). Main mesocarnivores are red fox (*Vulpes vulpes*), Eurasian badger (*Meles meles*) and wild cat (*Felix sylvestris*). Large carnivores besides grey wolf (*Canis lupus*) are Eurasian lynx (*Lynx lynx*) and occasionally brown bear (*Ursus arctos*) [42].

### Camera traps

Use of camera traps is a non-invasive, cost-efficient method for wildlife monitoring [46]. As it is a non-specific approach, one can study more species simultaneously [47] and also investigate human disturbance. The frequency of animals captured on a camera trap correlates with their abundance and activity and thus can be a useful indicator of the occurrence of species in a given location [48–50]. Furthermore, as indicated above, this method is also suitable for studying the daily activity pattern of species in a habitat [16,27,51,52].

We used RECONYX camera traps (UltraFire XR6; HyperFire HC500, PC900 and PC800; RapidFire PM75). They were posted on roads, trails, game trails, mud baths where the presence of the wolf was previously confirmed by tracks and scats. Cameras were inspected every 4 weeks to change batteries and memory cards. We analysed the data obtained from May 2015 to December 2019 (Fig 1) on twenty seven different sites. The recordings consist of both images and videos.

**Fig 1.**
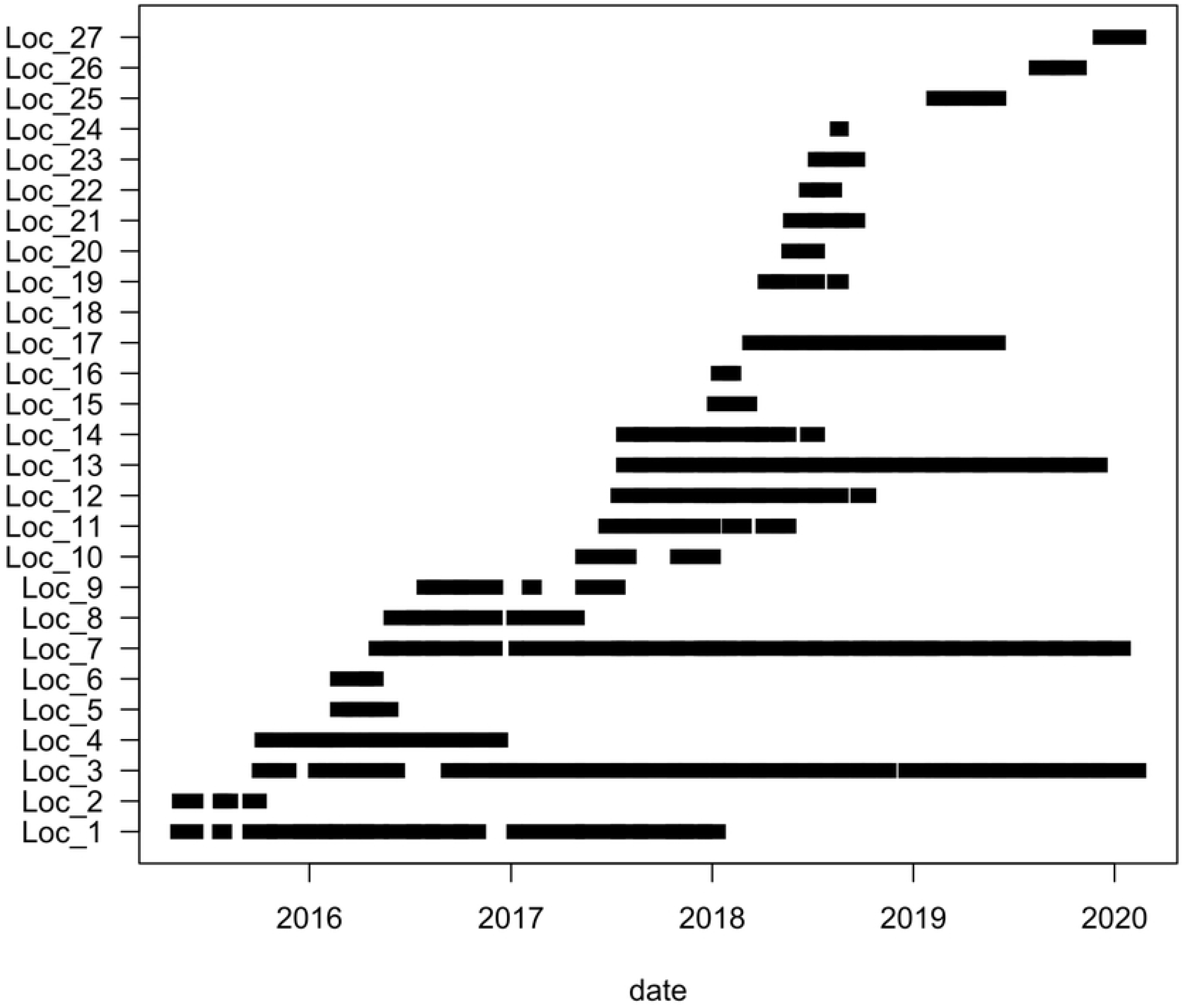
The operating periods of camera traps.

### The data

For each site, one of us (ZS) screened the recordings to collect the date and time of occurrences of studied species or human disturbance at a particular site. Following Ciuti et al. (2012) [11] we categorized human disturbance into three types: hikers (including hunters), motorised vehicles (m.vehicle) and riders (bikers and equestrians). The number of individuals (both animals and humans) were also collected. To lower the level of pseudoreplication we pooled two subsequent occurrences of the same species or the same type of human disturbance if they were separated by less than 15 minutes [15].

We analysed 101 676 files of 267.55 GB. We recorded 11 689 occurrences in total, of which were 596 *Canis lupus*, 1035 *Vulpes vulpes*, 1826 *Cervus elaphus*, 1738 *Sus scrofa*, 1301 *Capreolus capreolus* observations. Furthermore, 4139 observations of human disturbance, 1570 hikers, 2312 motorised vehicles, and 257 riders, were recorded. The remaining 1054 observations were excluded from the analyses because they included infrequently observed mammals, such as *Meles meles, Martes foina, Ovis gmelini, Sciurus vulgaris* and *Lepus europaeus*, livestock and indeterminable observations.

All analyses were performed in the R interactive statistical environment (version 4.0.4, R Core Team 2021).

### Temporal Overlap

We estimated temporal overlap between (i) games and wolf, (ii) human activity and games and (iii) wolf and human activity through the non-parametric calculation of the overlap coefficient (Δ_4_: Weitzman 1970). Δ_4_ ranges from 0 to 1, where 0 means no overlap, and 1 means complete overlap [53]. We classified site locations as low human density (LHD) areas where human activity is below the median human activity detection per day and high human density (HHD) areas otherwise. Separate temporal overlap estimations were performed on LHD and HHD classes, as well as over every site. Confidence intervals on Δ_4_ values were computed after bootstrapping samples 100 times from the dataset. We performed this analysis with the „circular” and „overlap” R packages [54,55]. We used two-sample t-tests to compare overlap coefficients between LHD and HHD locations.

### Co-occurrence of game and disturbance factors

To investigate the effects of predators (e.g. wolf) and human disturbance (e.g. motorised vehicles or hikers) on the occurrence of big game we fitted eXtreme Gradient Boosting (XGBoost) models as implemented in the ‘xgboost’ R package (version 1.6.0.1) [56]. To increase the robustness of our analyses we only considered presence/absence data, i.e. we re-coded each variable as one if the given species or human disturbance occurred on a given day and zero otherwise. As we are mainly interested in the relative effects of the presence of disturbance factors (predators or humans) we removed those days from the dataset when neither the game nor the disturbance factors occurred. In other words, we deleted those rows where all occurrence values were zero. This step prevents us from estimating the absolute probability of occurrence but it facilitates the model fitting process, e.g. by allowing the calculation of ten-fold cross-validation statistics.

For each of the three big games (*C. capreolus, C. elaphus* and *S. scrofa*) we run separate analyses. In these analyses the occurrence data of the given big game served as label or dependent variable, while the occurrence data of human disturbances (motorised vehicle, hikers, riders) and those of wolf (for all three big game) and fox (only for *C. capreolus*, as it is important predator for fawns [57]) were used as features or explanatory variables.

To infer the reliability of our results we run permutation tests where the occurrence data of big game (our dependent variables) were randomised within each stratus formed by each operating period within each camera. For both the observed values and the randomised datasets we calculated (i) metrics characterising the model fits and (ii) the differences in occurrence probability of the big game predicted for the presence and absence of the given disturbance factor. We used the following metrics: the square root of mean squared error (rmse), the logarithmic loss (logloss) and the area under the curve (auc) [56]. To assess reliability we compared the calculated metrics and differences to their distributions obtained from the randomised fits. Here after, we call an effect as significant if its value calculated from observed data is not contained by its distribution obtained from the randomised data. Each randomisation was repeated 1000 times.

All model fits were run for ten rounds with the default parameter values of the xgboost package.

## Results

### Temporal Overlap

Predators and games are mainly active during the night, while human activity peaks at the middle of the day (Fig 2). The Δ_4_ values between human and game activities are lower than values between wolf and games. In the case of games-human overlaps we found the greatest Δ_4_ value for roe deer (Δ_4_=0.52), followed by red deer (Δ_4_=0.41) and finally the wild boar (Δ_4_=0.26). The wolf has an overlap value with human activity (Δ_4_=0.40) which is comparable to the overlap values between humans and games. For the wolf-games overlaps the greatest overlap value was for red deer (Δ_4_=0.89) followed by roe deer (Δ_4_=0.84) and wild boar (Δ_4_=0.83). The 95% confidence intervals on the calculations are shown in Table 1. These indicate that the overlap between wolf and games are significantly higher than between human activity and games. Comparing the Δ_4_ values calculated separately for LHD and HHD areas, we can observe significant increases in overlap in case of human disturbance with all games at LHD sites. Contrary to this effect, wolf-game overlaps decrease significantly at LHD sites. In case of human-wolf overlap, the overlap also decreases significantly.

**Fig 2.**
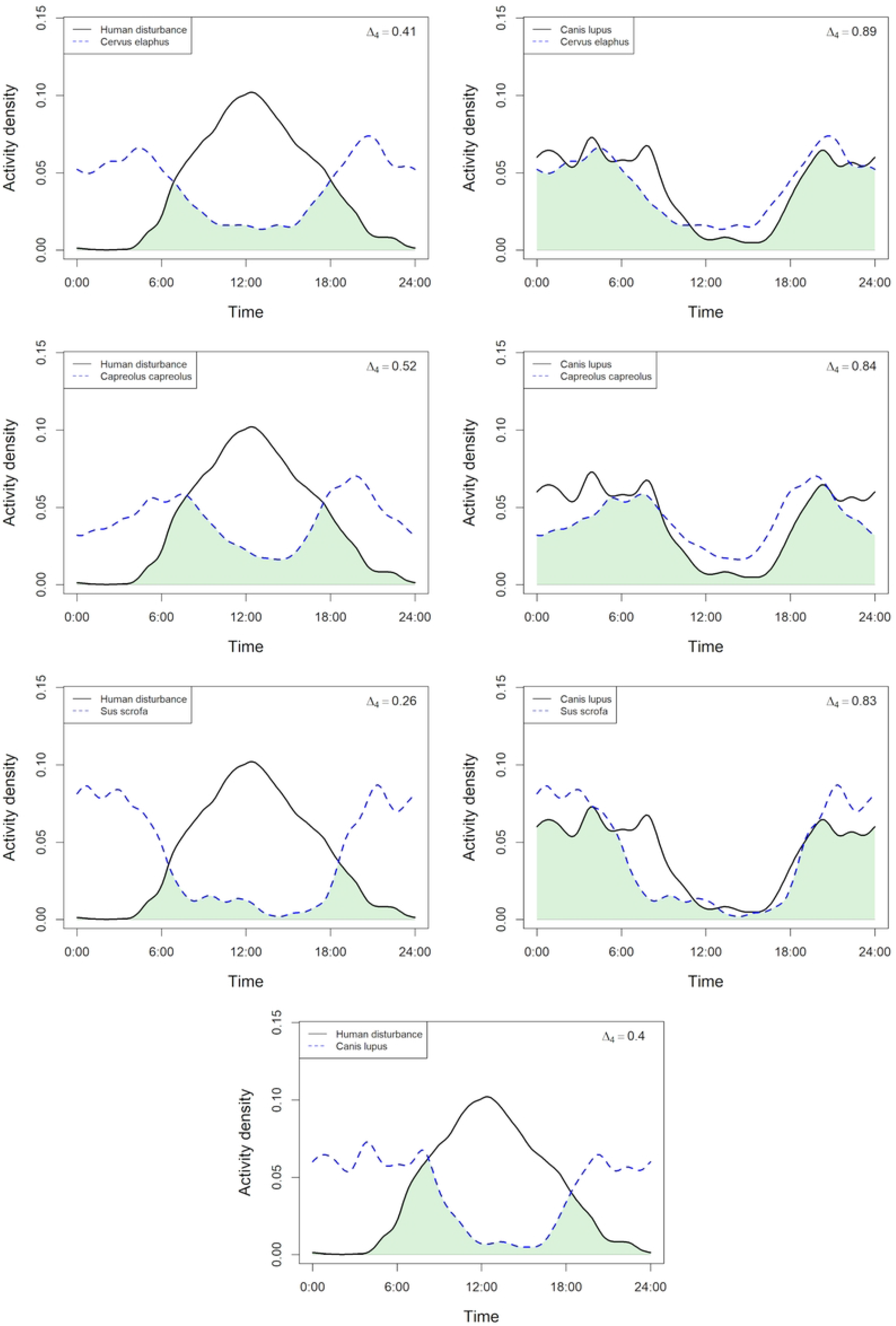
Temporal overlap between human activity and big games, between *Canis lupus* and big games, and between human activity and *Canis lupus* for all areas. The estimated overlap values (Δ_4_) between two activity patterns range from 0 to 1, where 0 means no overlap, and 1 means complete overlap.

**Table 1.** Summary of overlap coefficients (Δ_4_, for all sites), their 95% confidence intervals (95% CI), and comparison of Δ_4_ values between Low Human Density (LHD) and High Human Density (HHD) sites.

| Overlap | $\Delta_4$ | 95% CI | HHD $\Delta_4$ | LHD $\Delta_4$ | $\Delta_4$ change | p-value (FDR) |
| --- | --- | --- | --- | --- | --- | --- |
| Human with |  |  |  |  |  |  |
| *C. capreolus* | 0.52 | [0.49; 0.55] | 0.49 | 0.55 | +0.06 | p<0.001 |
| *C. elaphus* | 0.41 | [0.39; 0.43] | 0.37 | 0.47 | +0.10 | p<0.001 |
| *S. scrofa* | 0.26 | [0.24; 0.27] | 0.22 | 0.24 | +0.02 | p<0.001 |

| **Wolf with** |  |  |  |  |  |  |
| --- | --- | --- | --- | --- | --- | --- |
| *C. capreolus* | 0.84 | [0.80; 0.88] | 0.83 | 0.73 | -0.10 | p<0.001 |
| *C. elaphus* | 0.89 | [0.86; 0.92] | 0.88 | 0.75 | -0.13 | p<0.001 |
| *S. scrofa* | 0.83 | [0.79; 0.86] | 0.80 | 0.69 | -0.11 | p<0.001 |
| **Human with wolf** | 0.40 | [0.37; 0.43] | 0.40 | 0.33 | -0.07 | p<0.001 |

Significant differences were found between LHD $\Delta_4$ and HHD $\Delta_4$ values with $p < 0.001$ after false discovery rate (FDR) correction. $\Delta_4$ change was calculated as LHD $\Delta_4$ - HHD $\Delta_4$ .

### Co-occurrence of game and disturbance factors

For all three big games the metrics of the fits indicate that our models fitted rather well (Fig 3).

**Fig 3.**
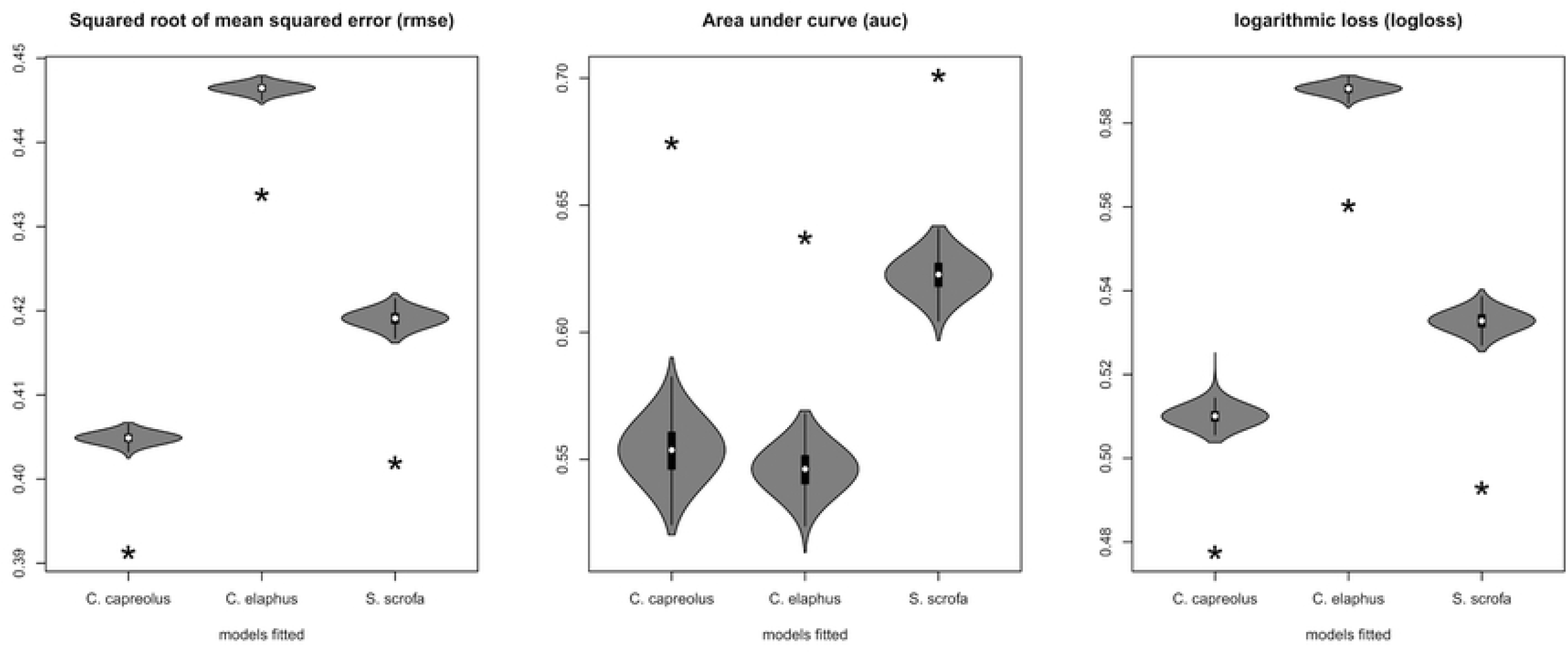
Metrics characterising the fit of XGBoost models to the occurrence data of big games in the Bükk Region. The stars show the metrics for the models fitted to the observed data. The violin plots illustrate the distribution of 1000 metrics obtained by fitting models to randomly permuted data. In the cases of the metrics of rmse and logloss lower values indicate better fit. In case of auc, higher values mean better fit.

Examining the importance values of the explanatory variables (predators and human disturbance) reveals that the most important factors influencing the occurrence of big games are motorised vehicles (0.29-0.46) and hikers (0.23-0.35) (Fig 4).

**Fig 4.**
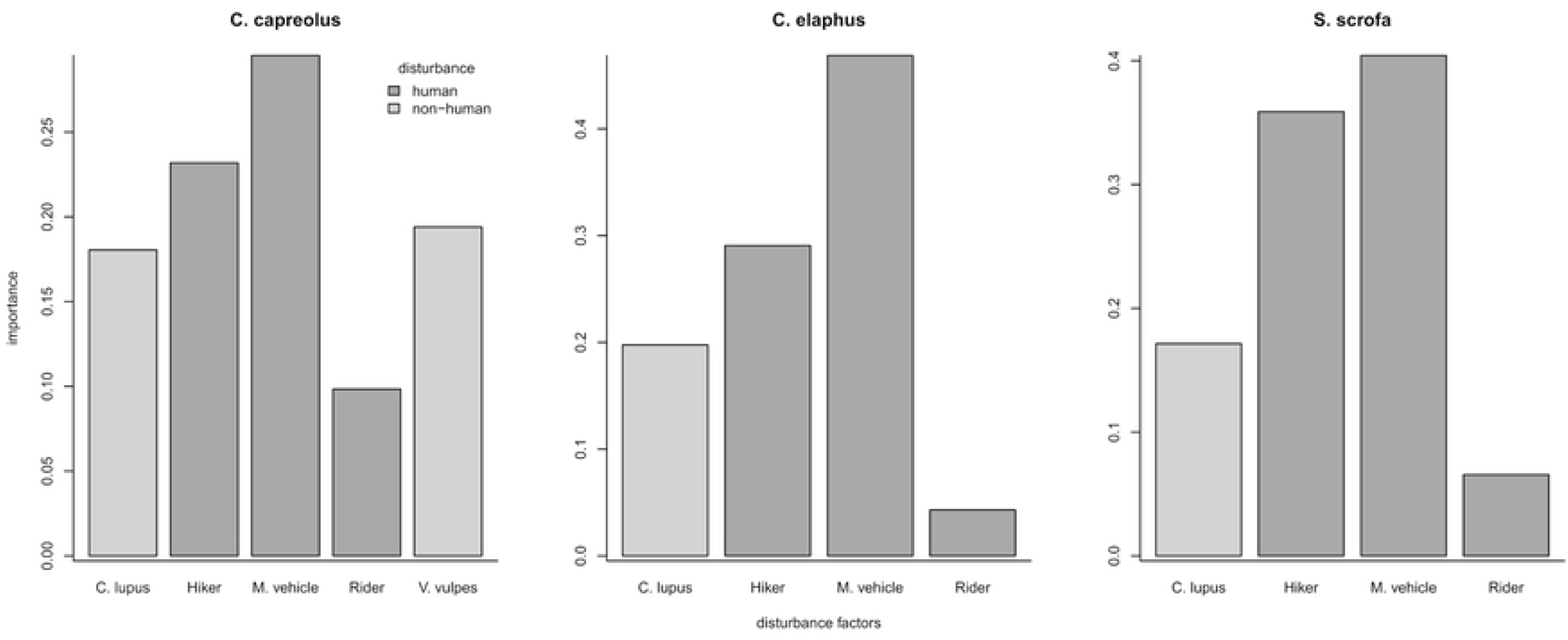
The importance of disturbance factors on the occurrence of big games in the Bükk region. Importance indicates how important a given factor is in the prediction of occurrence of big games. Darker columns indicate human disturbance, lighter ones predators.

Predators only appeared as third, in case of *C. capreolus* is fox (0.19), while in cases of *C. elaphus* and *S. scrofa* is wolf (0.19 and 0.17) and in case of *C. capreolus*, wolf (0.18) is forth. Riders were the least important factors in the cases of all three big games (0.04-0.09).

The effects of all disturbance factors were negative for each big game (Fig 5). In other words, the big games occurred less likely on the same day if either human disturbance or predators were present. The most significant of these were the effects of motorised vehicles and hikers while the effect of riders was never significant. In case of *C. capreolus* the influence of wolf wasn’t significant, because permuted data results were between -0.0042 and -0.0992, and observed data was -0.0952, also in case of *S. scrofa*, where permuted data was between -0.0438 and -0.1400, and observed data was -0.1406. Wolf influenced significantly in case of *C. elaphus*, where permuted data was between -0.0222 and -0.0855, while observed data was -0.1149 (min. difference is 0.0294). However human disturbance has a stronger effect in case of hikers (permuted data was between -0.0024 and -0.0712, observed data was -0.1333; min. difference is 0.0621) and motorised vehicles (permuted data was between -0.0169 and -0.0879, observed data was -0.1254; min. difference is 0.0375).

**Fig 5.**
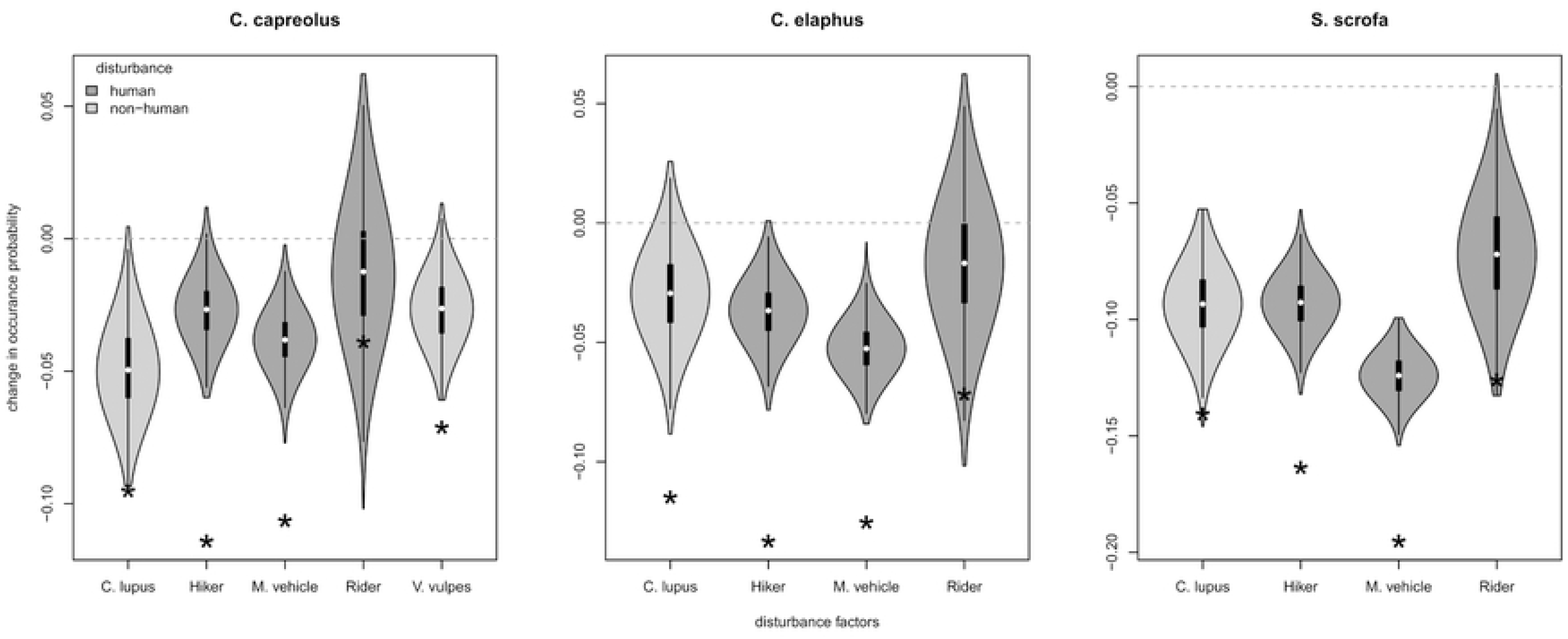
The estimated effects of disturbance on the occurrence of big games in the Bükk region. The stars mark the effects estimated from the observed data, while the violin plots illustrate the distributions of effects from randomly permuted data. Darker violins indicate human disturbance, lighter ones predators.

## Discussion

Games can adapt to predators and human disturbance by limiting their temporal or spatial overlap in their activities. According to previous observations, games often change their daily activity pattern if a predator appears in the same habitat [23–27]. Predators, however, are also able to adjust their activities to match the games’ [27,28].

Our results show that the activity patterns of animals and humans are markedly different. The games and wolf are mainly active at night and dawn, while human activity is predominantly observable at daytime and peaks at noon. Human activity can modify the activity patterns of both predators and games: the transition of diurnal animals to nocturnal life caused by human activity is a well-known, globally observed phenomenon [8,58]. Also human disturbance can change the predator’s activity patterns indirectly through the changes caused in activity patterns of games [28]. Our overlap analyses also indicate that games avoid humans more in areas where they have a higher chance to meet them (HHD areas). Furthermore, high human density results in higher overlap between wolves and games which might indicate that games accept a higher overlap with wolves to decrease the overlap with humans i.e. humans force the prey to overlap more their activities with predators. These results suggest that games actively avoid humans.

Temporal changes in activity patterns can have negative effects on the fitness of animals [8,58]. For instance, the chance of successful hunting and foraging may significantly decrease when a basically diurnal species is forced to become nocturnal which can ultimately result in the reduction of the success of mating and parental care. Furthermore, to games, the anti-predator behaviour may also be less efficient because of the poorer visibility [8,58]. Apart from temporal avoidance, humans can also influence the spatial distribution of games and predators [1,8,19,59]. By investigating land use differences Theuerkauf and Rouys (2008) [9] found that games did not avoid the areas used by wolves.

Based on our XGBoost results we could only detect a mild negative effect of wolves on the occurrence of red deer. This finding corroborates the results of Lanszki et al. 2012 [36], i.e. that red deer is the main prey of wolves in this area. Nevertheless, the presence of wolves was not the most important factor influencing game occurrence, the negative effects of human disturbance were larger and more important in the case of all games. As our analyses show, the presence of motorised vehicles, followed by hikers influenced most the occurrence of big game in this area. In accordance with results of Ciuti et al. (2012) [11] we also couldn’t detect significant effects of equestrians or bikers (riders).

Because of the return of the grey wolf the games are now subject to additional pressure. From an hunting economic point of view this is mainly important in the case of red deer and the wild boar since they are economically the most important big game species in the region. Our results, however, suggest that this additional pressure, compared to the strong human disturbance, is relatively mild for these species. Nevertheless, as other studies have showed, this pressure can facilitate the increase of the local biodiversity [1–4,10]. We also have to note that based on our results human disturbance has a larger influence on the behaviour of big game species than the presence of wolves.

To summarise, our results indicate that human disturbance has a strong influence for the game species. The disturbance by motorised vehicles is especially dominant. Therefore, intermittent resting in the woods, like restrictions on vehicle use, or reducing pedestrian traffic to some places might have an advantageous effect on the populations of the big game.

## Acknowledgements

We are grateful to the Bükk National Park Directorate, and their rangers, who gathered the data, especially to Péter Mlakár and Ádám Pongrácz. We thank Tamás Cserkész for his careful comments on a previous version of the manuscript. ZB was supported by a National Research, Development and Innovation Fund grant (TKP2021-NKTA-32).

